# Fluorescence-free quantification of protein/nucleic-acid binding through single-molecule *kinetic locking*

**DOI:** 10.1101/2020.09.30.321232

**Authors:** Martin Rieu, Jessica Valle-Orero, Bertrand Ducos, Jean-François Allemand, Vincent Croquette

## Abstract

Fluorescence-free micro-manipulation of nucleic acids (NA) allows the functional characterization of DNA/RNA processing proteins, without the interference of labels, but currently fails to detect and quantify their binding. To overcome this limitation, we developed a new method based on single-molecule force spectroscopy, called *kinetic locking*, that allows a direct *in vitro* visualization of protein binding while avoiding any kind of chemical disturbance of the protein’s natural function. We validate *kinetic locking* by measuring accurately the hybridization energy of ultrashort nucleotides (5,6,7 bases) and use it to measure the dynamical interactions of E. coli RecQ helicase with its DNA substrate.

---

Single-molecule (sm) micro-manipulation techniques such as AFM, optical tweezers or magnetic tweezers, have extensively been used to study NA. The approach relies on stretching NA by applying pico-Newton-scale forces and measuring their extension changes upon binding of the molecules of interest. While limited to *in vitro* measurements, they reveal features that are neither observable with bulk measurements nor with *in cellulo* fluorescence microscopy. For example, they unveiled the precise mechanisms of NA unwinding and replication by enzymes such as helicases and polymerases [1]. However, the individual binding events of proteins with NA can not be directly observed because they generally do not imply sufficiently large extension changes. Spatiotemporal resolution of single-molecule techniques is still making progress [2] and now allows observing the nanometric signals created by oligo hybridization (>=8 nucleotides (nt)) [3] and helicase stepping [4]. However, the detection of simple binding events that involve Angström-scale extension changes below the second timescale are still out of reach. Coupling micromanipulation with Förster resonance energy transfer (FRET) fluorescence alleviates these lacks in spatiotemporal resolution [5]. Nevertheless, it is subject to standard fluorescence limitations, the most cumbersome being the need of substrate labeling and the uncertainty concerning the perturbation of the measurements by the fluorescent probe [6].

We report here a detection scheme for single-molecule micro-manipulation, called *kinetic locking*, that is neither based on extension changes nor on fluorescence. Detection is performed through the observation of the dynamics of a DNA fluctuating probe that is sensitive to changes in its direct chemical environment. The probe consists of a 10-bp DNA hairpin stretched at a force of 10 pN. At this force, the hairpin fluctuates between its duplex (closed) and single-stranded (ss, open) states [7] (Figure 1.A). The typical times between two successive changes of state is 10 ms (Figure 1.B). These fluctuations are highly sensitive to the applied force (a force increase of 0.7 pN doubles the opening rate of our probe), to the sequence of the hairpin [8], its apex loop length, and the presence of mismatches [9]. We also noticed that the kinetics is highly altered by the binding of another molecule. For example, a protein binding to ssDNA prevents the closing of the hairpin and will thus increase the time spent in this opened state. On the other hand, a protein binding to dsDNA prevents the opening of the hairpin and will consequently increase the time spent in its closed state. Therefore, these altered fluctuations can be used to detect binding events. In order to constitute a sensitive probe, the transition rates of the hairpin must however display a strong temporal stability. We thus chose magnetic tweezers over AFM or optical tweezers [10] given their unrivalled stability in force and in temperature. Our setup allows us to keep the relative variation of the transition rates (typically 100 s^−1^) of the probe below 5% during several hours (Supplementary Figure S1).

**Figure 1:**
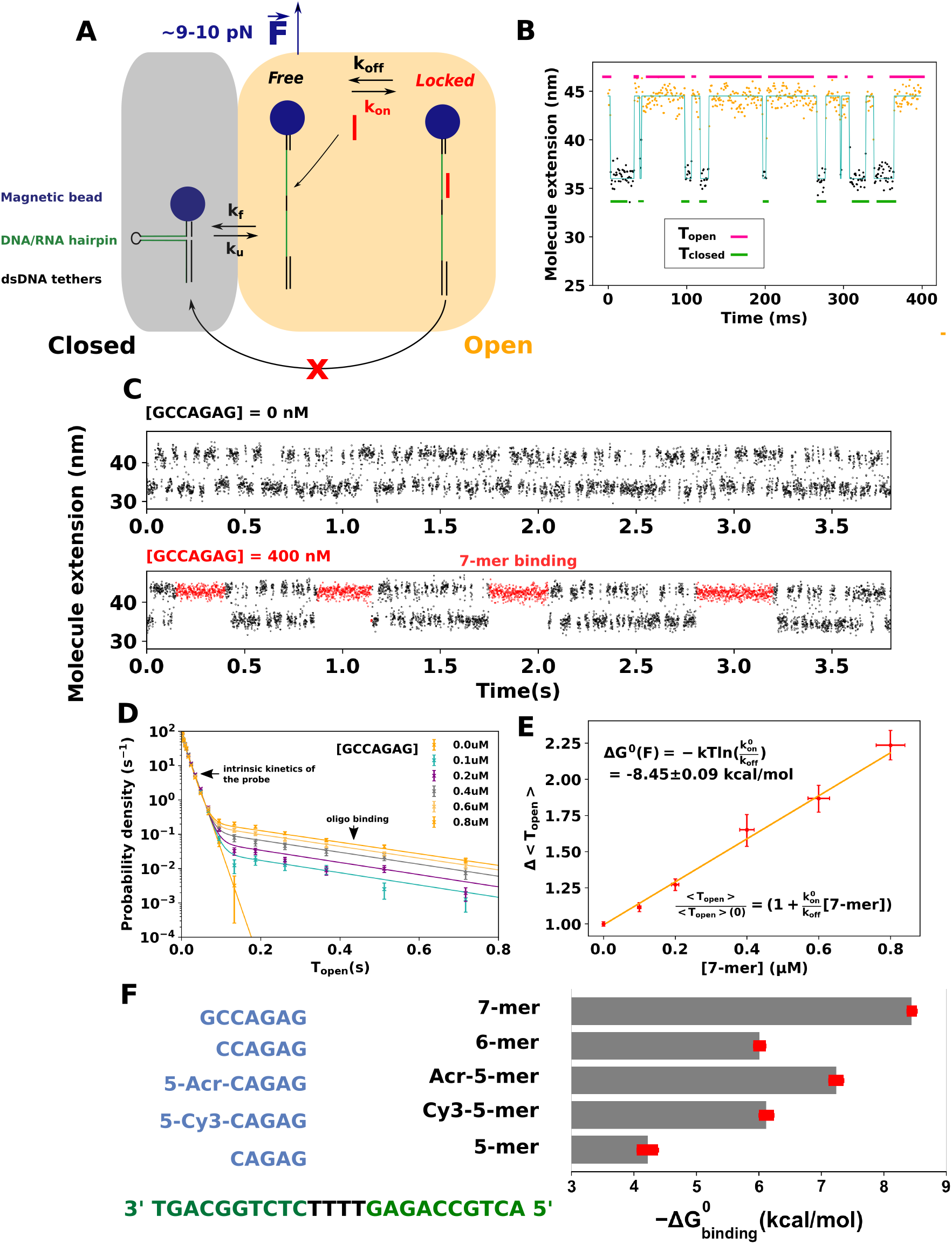
Measurement of the hybridation energy of short oligonucleotides with kinetic locking. **A.** Description of the assay. When an oligonucleotide binds to the probe, it blocks it transiently in the open state. **B.** Measured extension of the hairpin as a function of time. Dots represent raw data. The blue line and the dot colors represent the inferred state. T_open_ and T_closed_ are defined as the time spent in the open and closed states by the hairpin. **C.** Larger time view of the measured extension of the hairpin in the absence of oligonucleotide in solution (top) and in the presence of the complementary 7-mer (bottom). **D.** Distribution of the times spent by the DNA probe in its open state as a function of the concentration of 7-mer. The clear double exponential distribution allows to deduce the binding rates k_on_ and k_off_. **E.** Relative evolution of the mean time spent by the DNA probe in the open state <T_open_> as a function of the concentration of 7-mer. The slope allows to compute the energy of binding. Y-errors are computed through bootstrap resampling. X-errors are based on a putative error of 5% on the concentration. Each data point consists in the averaging of at least 7000 events. **F.** Free energies of binding of five different oligonucleotides measured with kinetic locking. The sequence of the probe is shown in green and the sequences of the oligonucleotides in blue. Cy3 and acridine modifications increase substantially the binding energy. Errors are based on the covariance of the parameters inferred from the linear fits as the one shown in E.

First, we show how *kinetic locking* enables the direct visualization of the binding of a 7-bases oligonucleotide. When the fluctuating DNA hairpin is stretched at 10 pN and no oligonucleotide is present in solution, the distribution of the times *T*_open/close_ spent in its closed/open states follow clear single-exponential distributions of parameters *k*_*f*_ (folding rate) and *k*_*u*_ (unfolding rate). Typical opening/closing times lie in the 10ms range. Upon the injection of an oligonucleotide complementary to the hairpin sequence, the hairpin stays locked in the open state during ~250 ms (Figure 1.C, bottom). We interpret this time at the typical binding time of the oligonucleotide. This is confirmed by the change of the distribution of the open times. The latter goes indeed from single to double-exponential (Figure 1.D), as predicted by the model underlying *kinetic locking* (Supplementary S.I), and allows us to calculate the kinetic rates of binding of the 7-mer at ~9.5 pN (Table S1).

Next, we validate the accuracy of the method by performing the first single-molecule measurement of the thermodynamics of the hybridization of very short oligonucleotides (5, 6 and 7 bases). Indeed, our model predicts that the average time spent in the open state by the fluctuating probe should increase linearly with the concentration of oligonucleotides, the slope depending on the free energy of hybridization Δ*G*_binding_.

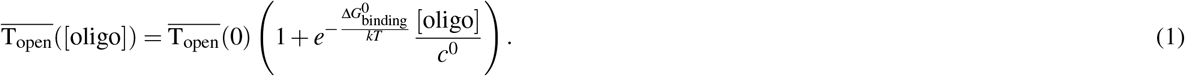

Interestingly, this dependence is also valid for the fast hybridization of 5-mer and 6-mers oligonucleotides whose typical association and dissociation times are smaller (< 10 ms) than the typical fluctuating times of the probe. Our experiments confirm the linear dependence of 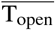 with the concentration of oligomers and the absence of dependence of the times spent in the closed state 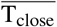 (Figures 1.E, S2 and S3). To control that the effect is not due to an increase of ionic strength due to the presence of charged NA, we show that introducing an oligonucleotide that is not complementary to the sequence of the fluctuating hairpin (C-C mismatch in third position) results in the absence of measurable change (Figure S2, second panel). We compare in Table 1 the ΔG obtained with our measurements with the widely used nearest-neighbor (NN) tables [11]. Usual salt corrections were used [12, 13]. Our results are in very good agreement with the data, thus validating our method. We also quantify the effect of adding an acridine group and a Cy3 group at the 5’ position of the 5-mer. The modified oligonucleotides display a larger or equal hybridization energy than the 6-mer, confirming their strong interaction with DNA (Figure 1.F). So far, temperature-jump IR spectroscopy is the only technique that was able to provide data [14] on the thermodynamics of ultrashort nucleotides. However, its accuracy is limited by the deuterium effects on hydrogen bonding and the short rethermalization time *T*_th_ : in order to keep the association times smaller than *T*_th_, unrealistic salt conditions must be used ([Na] ~200mM and [Mg] ~40mM). In addition to validate the method, the measurement of the hybridization thermodynamics of such short oligonucleotides is thus interesting *per se*. It will help understanding the role of specific sequence motives on biological processes that involve transient hybridization such as silencing through RNA interference and to better understand unwanted off-target effects [15]. It will also help identifying chemical modifications that stabilize short duplexes strongly enough so as to allow mechanical sequencing [16].

**Table 1.**
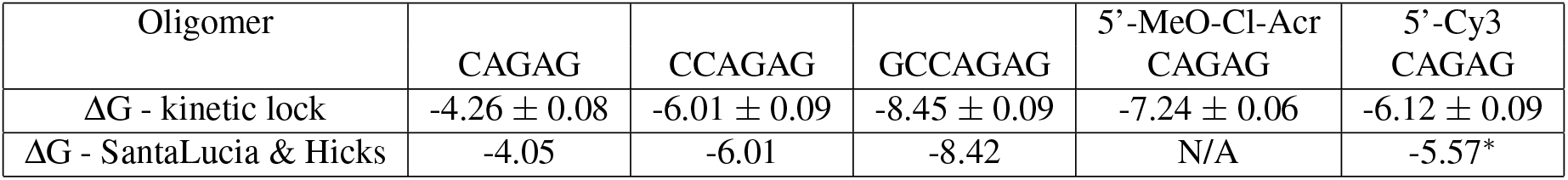
Comparison of the experimental binding energies (in kcal/mol) obtained by *kinetic locking* with standard predictions by the nearest-neighbor model at 25°C. NN-computations were made with the freely available software Biopython [17] after correction of two misreproduced values from the cited thermodynamic tables. Displayed errors are statistical errors. The application of a force of 9.5 pN induces a systematic correction that lies between −0.015 and 0.040 kcal/mol/nucleotide (See Supplementary S.V). All curve used to infer these values are shown in Supplementary Figure S2. ^∗^The effect of the Cy3 group is estimated using the sequence dependent estimations provided by Moreira *et al.* in [6].

| Oligomer | CAGAG | CCAGAG | GCCAGAG | 5'-MeO-Cl-Acr<br>CAGAG | 5'-Cy3<br>CAGAG |
| --- | --- | --- | --- | --- | --- |
| $\Delta G$ - kinetic lock | $-4.26 \pm 0.08$ | $-6.01 \pm 0.09$ | $-8.45 \pm 0.09$ | $-7.24 \pm 0.06$ | $-6.12 \pm 0.09$ |
| $\Delta G$ - SantaLucia & Hicks | -4.05 | -6.01 | -8.42 | N/A | -5.57* |

We finally use the *kinetic locking* scheme to investigate the DNA-binding kinetics of the catalytic core of the E. coli helicase RecQ, RecQ-ΔC (Figure 1.A of [18]), in the absence of free nucleotide in solution. Figure 2 shows the observation of the binding of RecQ-ΔC with DNA. The first probe (Figure 2.A, left panel) does not contain any single-stranded gap when closed. Due to its poor affinity to double-standed DNA [19], RecQ-Δ*C* can thus only bind in the open state, preventing the closing of the probe (Figure 2.A, middle panel). The distribution of times spent in the open state by the probe (Figure A, right panel) allows us to determine the binding parameters of RecQ-ΔC in this configuration : 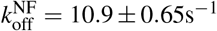 and 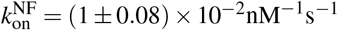. Furthermore, RecQ is known to display a stronger affinity for substrates containing a replication fork with a gap on the leading strand (LeGF) [19]. In order to assess the associated binding rates, we designed a slightly different fluctuating probe that contains a 14-nt single-stranded gap on the leading strand in its closed state (Figure 2.B, left panel). In addition to blocking the substrate in its open state, RecQ also binds to the closed hairpin and maintains it in the closed state (Figure 2.B, middle panel and Figure 2.C). These binding events are longer, clearly distinguishable, and their frequency increases linearly with the concentration of RecQ-ΔC in solution (Figure 2.D). We found 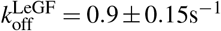 and 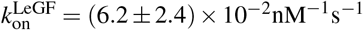. Figure 2.E shows the 10-fold difference of dissociation rates between the configuration containing a fork with a leading gap (LeGF) and the configuration that does not (NF). *k*_off_ is smaller in the case of the forked-substrate. This confirms the larger affinity of RecQ for a forked substrate observed in bulk [19] with gel mobility shift assays, and allows us to quantify the associated binding rates. The fact that RecQ maintains the hairpin in its closed state corroborates structural studies showing that the cleft formed by the winged-helix (WH) and the Zn^2+^ finger wraps around dsDNA [20, 21]. It furthermore indicates that this cleft is strong enough to stabilize the fork hybridization while a force is applied. We hypothesize that this stabilization mechanism of fraying hairpin is used by helicases of the RecQ family helping polymerases to reload [22] and allowing the restart of stalled replication forks [23]. Interestingly, RecQ-ΔC also keeps the fork closed in the presence of a 7-bases gap on the nascent lagging strand (Figure S4), indicating that fork gripping by the enzyme is not dependent on the interaction with the free 5’-end.

**Figure 2:**
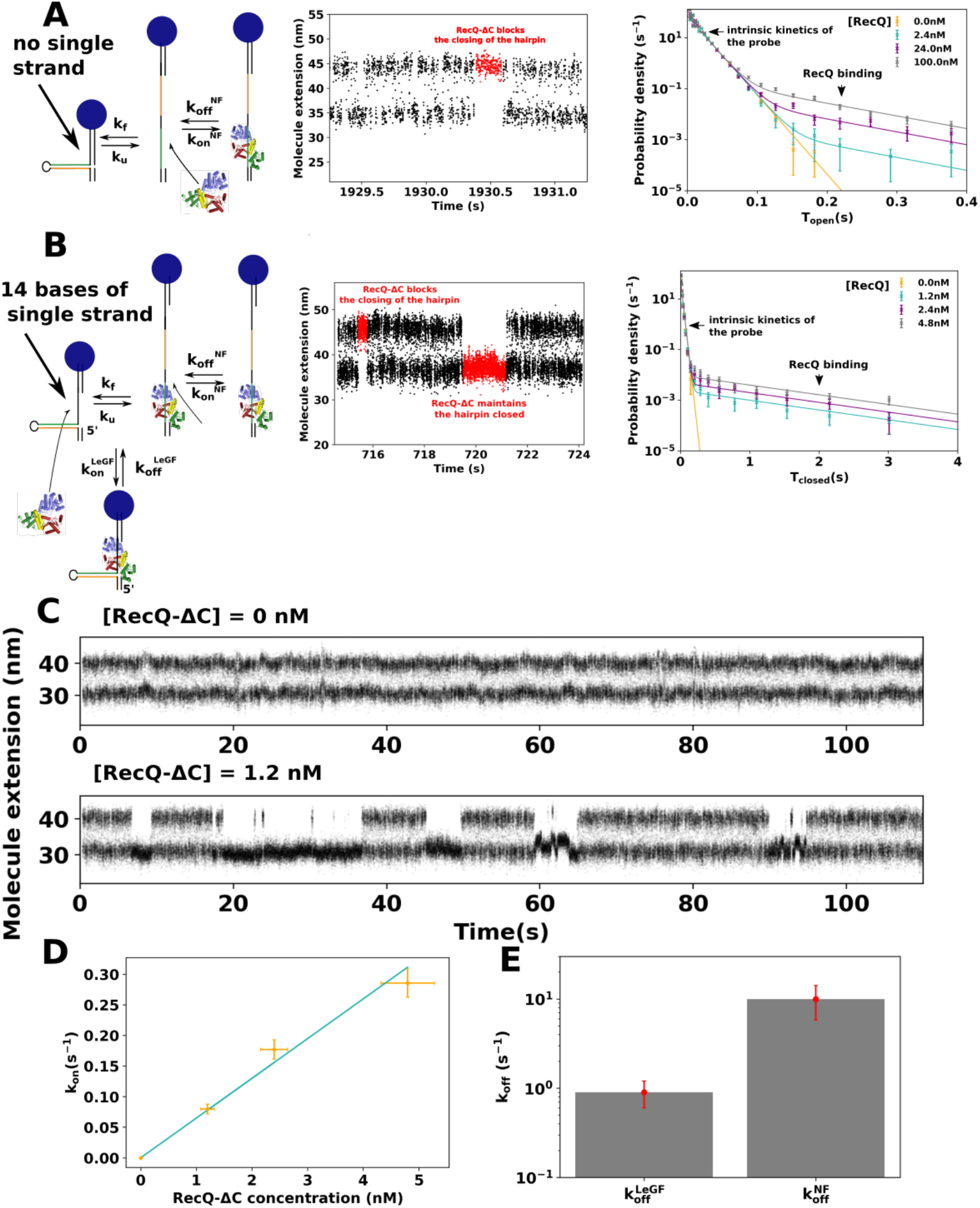
Detection and quantification of the binding of RecQ-ΔC with various DNA subtrates using kinetic locking. **A.** 10-bp hairpin without ssDNA gap (Left) is used as a fluctuating probe. (Middle) Binding of RecQ-ΔC can be observed through the transient blocking of the hairpin (HP) in its open state. (Right) Distribution of times spent by the hairpin in its open state as a function of RecQ-ΔC concentration, fitted by the double exponential law predicted by the kinetic model (Supplementary SI). Relative errors of a given bin are taken as 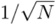, where N is the number of points in the bin. **B.** 10-bp hairpin with a 14-nt ssDNA gap (Left) is used as a fluctuating probe, simulating a leading-gapped replication fork, LeGF. (Middle) Binding of RecQ-ΔC on the ssDNA gap results in the transient stabilization of the hybridized state of the HP. (Right) Distribution of times spent by the hairpin in its closed state as a function of RecQ-ΔC concentration, fitted by a double exponential law. **C.** Larger time view of probe B. with and without RecQ-ΔC. **D.** Binding rate k_on_ in the closed state for configuration B. as a function of RecQ-ΔC concentration. Y relative errors correspond to the inverse of the squareroots of the number of observed events. X-relative errors are based on a 10% relative error on the concentration. **E.** Comparison of the dissociation rates k_off_ of RecQ-ΔC with (k^LeGF^) and without (k^NF^) the presence of a ssDNA gap. Error bars are derived from bootstrap resampling (see Methods).

In conclusion, the method that we report here allows the fluorescence-free detection of discrete protein/NA binding events, alleviating the need of protein labeling while avoiding the disturbance of the measured interaction by an external probe. The time-resolution presented here (~10 ms) largely exceeds label-free sm-methods based on periodic force changes [24, 25] that are well suited to study long dissociation times but not short and repetitive bindings. Besides, it could be further increased by using a high-speed camera to track the molecule extension (10 kHz) while concomitantly increase the folding rate of the probe by slightly decreasing the applied force. Compared to stopped-flow bulk assays, such as the one based on tryptophan fluorescence [26], it avoids photo-bleaching, hidden binding events, and the need to introduce chemical competitors to evaluate dissociation kinetics. More importantly, the single-molecule nature of the experiment gives important insights about the nature of the interaction between the protein and its substrate, as was shown by the unexpected stabilization of the probe by RecQ catalytic core. The main limitation is the narrow force range (8-15 pN), since the NA substrate needs to be stretched in order to induce its fluctuations. However, this corresponds to the typical forces felt by NA during transcription, translation or replication and is thus consistent with physiological conditions. Our assay can be applied to a broad spectrum of DNA/RNA-binding proteins (notably transcription factors, polymerases, helicases, primases), and used to study their sequence and substrate specificity. An other potential application is the precise quantification of the inhibition of enzyme/NA binding by potential antiviral drugs, in a context where replicative enzymes are increasingly considered as relevant targets (helicase-primase of Herpes [27], helicase nsp13 of SARS-CoV-2 [28]). By providing reliable single molecule measurements while avoiding the functional interference of labeling groups, *kinetic locking* will complement the toolbox of enzymologists for the kinetic characterization of NA/protein complexes.

## Methods

### Design of stable fluctuating probe

The fluctuating probe consists of a 10-bp DNA hairpin that rapidly oscillates between its open and close states. Using the high-precision optical setup described in [3], we measure the transition rates of the 10-bp hairpin by tracking the 10-nm difference of extension between its two states. The time between two transitions is of the order of 10 ms, allowing us to measure *N* = 10^4^ transitions during a typical acquisition time of 100s and thus to reduce the relative error made on the estimation of the kinetic rates to 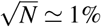. Supplementary Figure S1 shows the temporal stability of these rates. 7000s of acquisition are separated into 20 chunks of 350s. The kinetic rates inferred from each of them are all equal within a tolerance of 1 ms.

### Bead preparation

All single-molecule experiments are performed at 25°C. All the DNA substrate used in the presented assays are synthesized single stranded oligonucleotides. Their sequences are indicated in Table S2. Their 5’end is complementary to a 57 bases 3’ DBCO modified long oligonucleotide (Oli1) that is attached to azide-functionalized surfaces (PolyAn 2D Azide) through a 2-hour-long incubation (100nM Oli1, 500 mM NaCl). Their 3’end is complementary to a 58-base oligonucleotide (Oli2). Oli2 contains two biotin modifications at its 5’end. The ssDNA substrate is first hybridized with Oli2 by mixing both oligos at 100nM in 100 mM Nacl, 30 mM, Tris-HCl pH 7.6. 5uL of streptatividin coated Dynabeads MyOne T1 (Thermofisher) are washed three times in 200 uL of passivation buffer (140 mM NaCl, 3 mM KCl, 10 mM Na2HPO4, 1.76 mM KH2PO4, BSA 2%, Pluronic F-127 2%, 5mM EDTA, 10 mM NaN3, pH 7.4). The result of the hybridization between the substrate and Oli2 is diluted down to 2 pM buffer and then incubated 10 minutes with the beads in a total volume of 20uL of passivation buffer. The beads are then rinsed three times with passivation buffer in order to remove unbound DNA. 1uL of the bead solution is then introduced in the cell coated with Oli1 and filled with passivation buffer, and incubated for 5 minutes. Excess unbound beads are washed out by flowing passivation buffer.

### Sample preparation

For the DNA probes, three different constructs (HP1/HP2/HP3) are used, which sequences are given in Table S2. They consist of synthesized ssDNA of length respectively 160,153 and 139 bases. Once hybridized to Oli1 and Oli2, the size of the remaining single-stranded DNA reduces to (45/38/24) bases. 24 of these bases fold into a 10 bp hairpin with a 4-base apex T-loop.

The fluctuating probe used to measure oligonucleotide binding is HP1. Once the hairpin is hybridized, 7 ss bases are left on the 5’ side and 14 on the 3’ side. Measurements are performed in hybridization buffer (20 mM Tris-Hcl pH 7.6, 3mM MgCl2, 100 mM KCl). The oligonucleotides are ordered dry (Eurogentec) and diluted in the same buffer preparation as the one used during the hybridization assay in order to avoid any change in salinity upon injection. Concentrations are calculated based on the quantity indicated by the provider.

RecQ-ΔC is purified as described in [18] and is diluted without ATP in RecQ Buffer at a concentration specified in the main text. The fluctuating probes HP2 (with 3’ ssDNA gap) and HP3 (without 3’ ssDNA gap) are used. Measurements are performed in RecQ buffer (20 mM Tris-Hcl pH 7.6, 3mM MgCl2, 50 mM NaCl). The concentration of the enzyme is assessed through UV absorbance (210-340 nm) with a Nanodrop (Thermofisher) spectrophotometer (*ε* = 45840M^−1^cm^−1^).

### Data Acquisition

The extension of the fluctuating probe is acquired in real-time using the high-resolution magnetic tweezers setup described in [3] at an acquisition frequency of 1300Hz with a CMOS camera (UI-3030CP-M, IDS Ueye). The home-made C/C++ acquisition program *Xvin* can be found at https://tig.phys.ens.fr/ABCDLab/xvin/.

### Data analysis

The changes of states (open/close) of the fluctuating probe are detected automatically with the program *Xvin*, when the summed change of extension over a monotonous section of the extension curve is larger than 8 nm.

Times spent in the open state T_open_ correspond to the difference between the detected times of two successive opening and closing events. Times spent in the closed state T_closed_ correspond to the difference between the detected times of two successive closing and opening events.

Average times 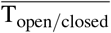 correspond to naive mean values. Errors are computed using bootstrap resampling. All averages of the hairpin closing and opening times are performed on samples containing at least 6000 independent events.

In the case of concentration-dependent double-exponential distributions, the times are fitted to the exact formula provided in Supplementary S.I with the SciPy [29] curvefit function. A single fit is used for all concentrations. Errors on the fit are computed based on bootstrap resampling : 100 samples of the same size as the original data sample are generated by resampling the original data with replacement. The fitting procedure is then applied to all samples, returning a distribution of inferred kinetic parameters. The values shown in the article correspond to the averages of these distributions while the errors correspond to their standard deviations. The proportion of long times in the double-exponential distribution is then verified with a binning-free expectation maximization (EM) algorithm based on the Pomegranate [30] Python module. All data analysis procedures can be found at https://github.com/Mriv31/kineticlocking.

## Author contributions statement

M.R. conceived, performed and analyzed the experiments. M.R. wrote the manuscript. M.R. V.C. and J.V-O. discussed the results. B.D. purified RecQ-ΔC. V.C. and J-F.A. built the magnetic tweezers and supervised the research. All authors reviewed the manuscript.

## Supplementary information

**Figure S1.**
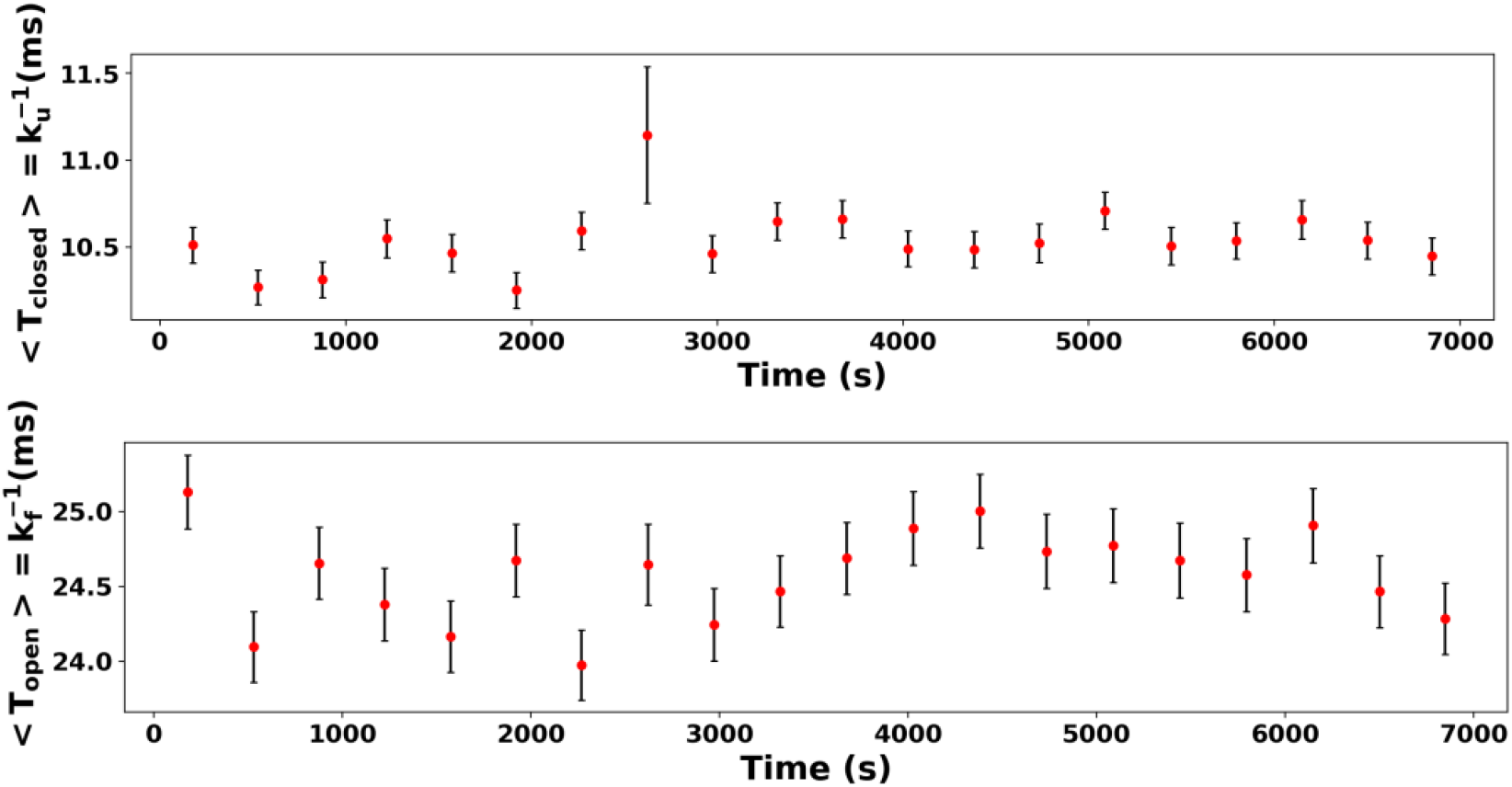
Average times spent in the closed and open states by the probe. Times are measured during 7000s of acquisition of the extension a single molecule. The average is performed over time bins of 350s. Error bars are calculated through bootstrap resampling.

**Table S1.**
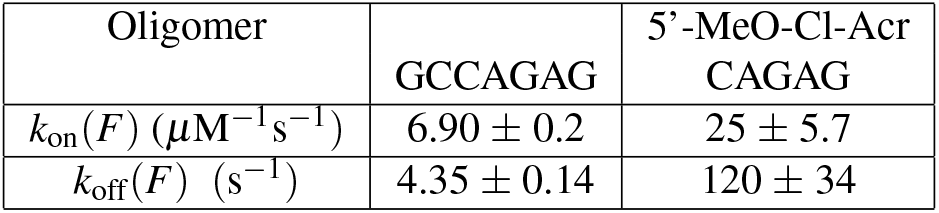
Binding (*k*_on_) and dissociation rates (*k*_off_) measured from *kinetic locking* at T = 25°C and F = 9.5 pN. Corresponding distributions from which they are inferred are shown in Figure S5

**Figure S2.**
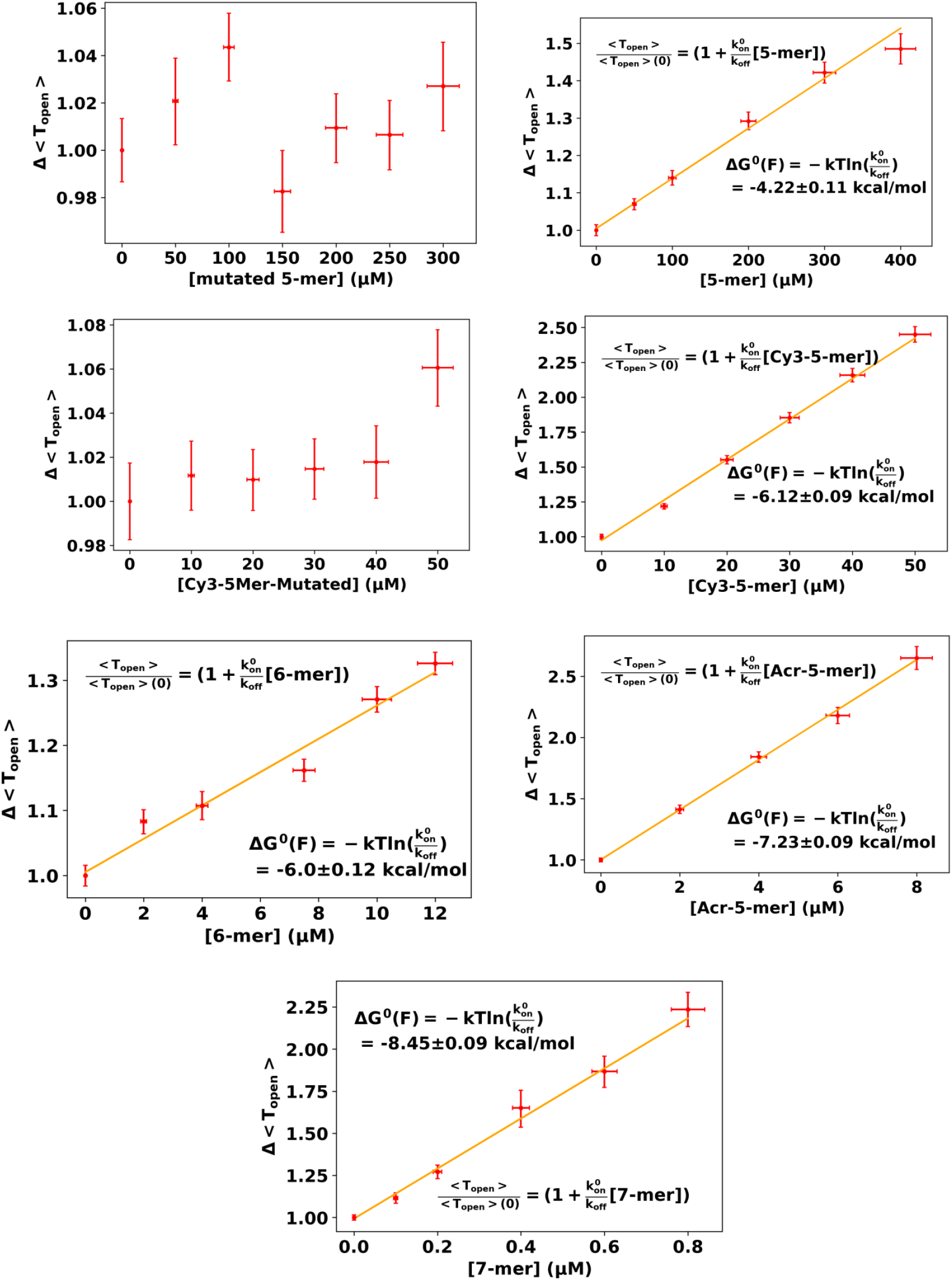
Average times spent by the fluctuating probe in its open state < *T*_open_ > as a function of oligonucleotide nature and concentration. Error bars are computed by bootstrap resampling. Errors on the fit are computed using the covariance matrix. Averages are performed on samples containing at least 6000 independent times. Mutated 5-mers (CACAG) have a mismatch at the third position compared to the oligonucleotides (CAGAG) that are complementary to the hairpin sequence.

**Figure S3.**
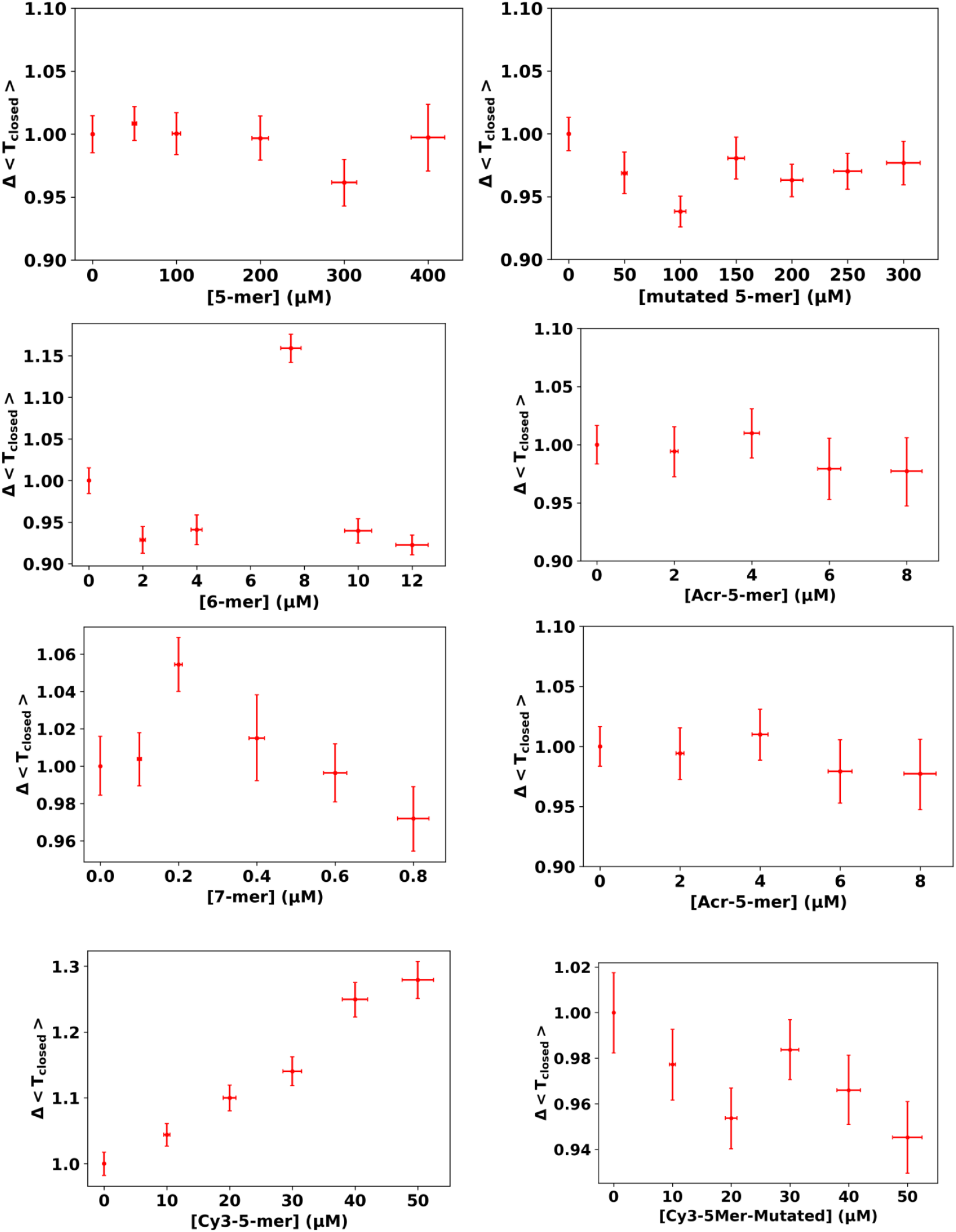
Dependence of the mean times spent by the fluctuating probe in its closed state as a function of oligonucleotide concentration. No significant change can be observed, with the notable exception of the 5-mer with 5’ Cy3 modification, that stabilizes the closed duplex state. Error bars are computed by bootstrap resampling. Averages are performed on samples containing at least 6000 independent times. Mutated 5-mers (CACAG) have a mismatch at the third position compared to the oligonucleotides (CAGAG) that are complementary to the hairpin sequence.

**Figure S4.**
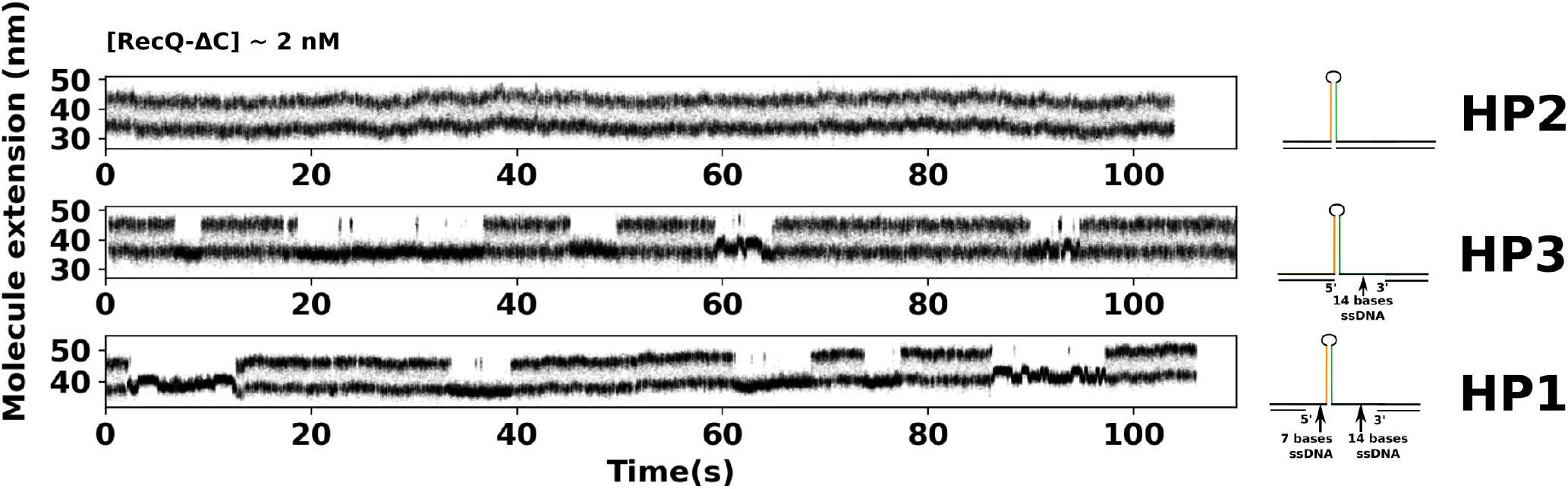
Time-course experiments showing the extension changes of the fluctuating probe in presence of RecQ-ΔC at 2 ± 1 nM for different probes (right) whose sequence can be found in Table S2. The stabilization of the closed state only happen when there is a single-stranded gap on the leading strand (HP1 and HP3), and is happening regardless of the presence of a single-stranded gap on the lagging strand (HP1).

**Figure S5.**
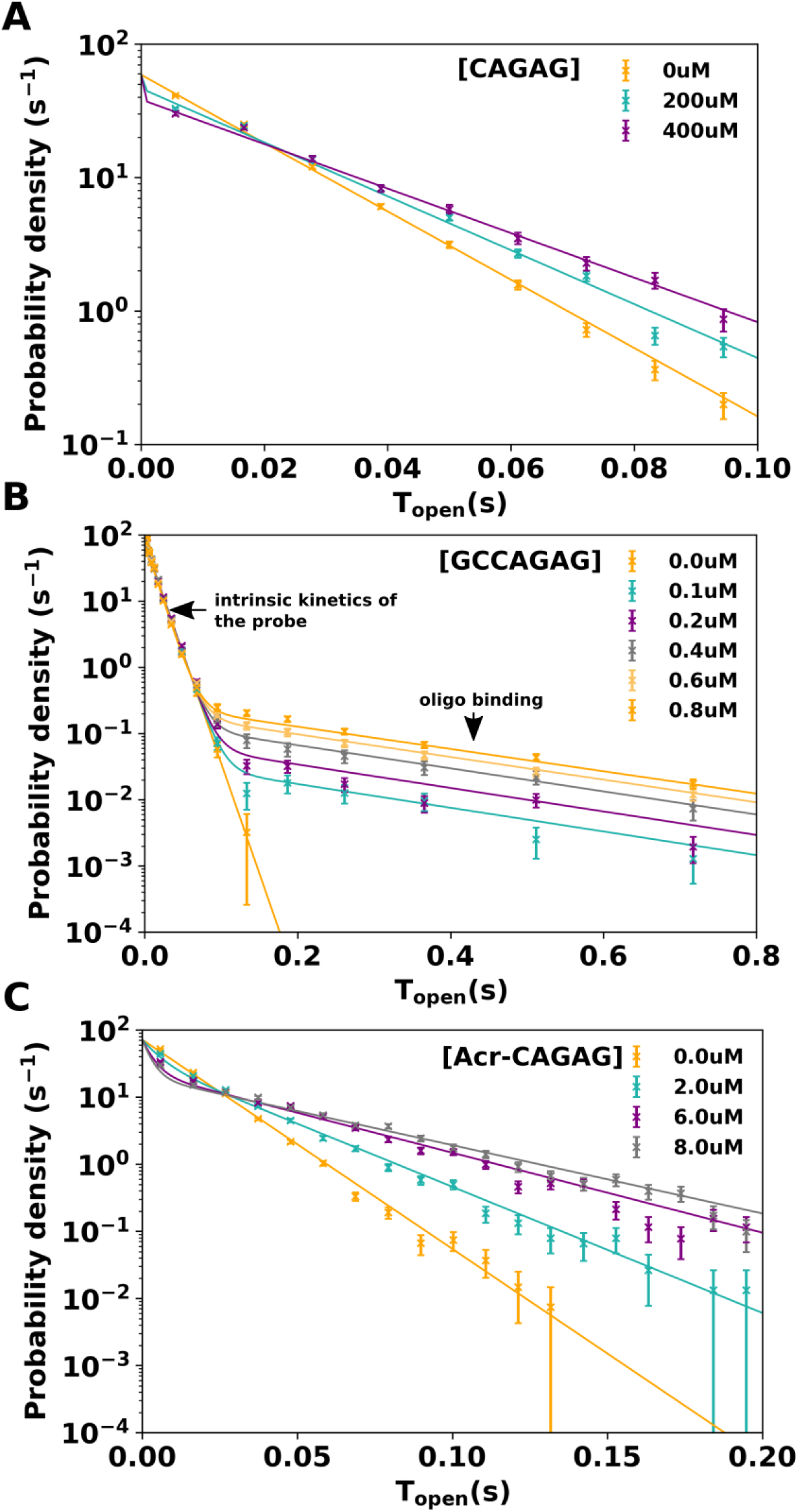
Distribution of the times spent by the DNA probe in its open state as a function of the concentration of oligonucleotides. For all oligonucleotides, a unique fit to the expected distribution (Supplementary SI) is performed. All concentrations are fitted together. **A**. 5-mer CAGAG. The kinetics of hybridization of the oligo being much faster than the kinetics of the probe, all distributions follow single exponential laws. *k*_on_ and *k*_off_ can not be inferred. (Supplementary SIII). **B**. 7-mer GCCAGAG. The kinetics of hybridization of the oligo being much slower than the kinetics of the probe, all distributions follow double exponential laws. *k*_on_ and *k*_off_ can be inferred. (Supplementary SII). **C**. 5-mer CAGAG with 5’ acridine modification. Intermediate case. *k*_on_ and *k*_off_ can be inferred but with a smaller precision due to the small proportion of short times.

### S.I Resolution of the model supporting *kinetic locking*

We compute the exact time distribution expected for a model of *kinetic locking* that follows the scheme described in Figure 1.A.

Calling *P*_*f*_ (*t*) the probability for the hairpin to be in the open free state (without oligo), *P*_*l*_ (*t*) the probability to be in the in the open locked state (oligo hybridized), and *P*_*c*_(*t*) the probability to be in the closed state, the systems follows the dynamics :

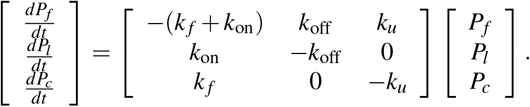

The free (*f*) and locked (*l*) open states are indistinguishable. The probability to be in one of these open states is called *P*_*o*_(*t*) = *P*_*f*_ (*t*) + *P*_*l*_ (*t*). Given a hairpin that just moved to the open free state (*P*_*f*_ (0) = 1, *P*_*l*_ (0) = 0, *P*_*c*_(0) = 0), we are interested in *ρ*_*o*_(*T*), the distribution of times spent in the open states before a closing event (*l* → *c*) happens. Such a closing event being observable, we are not interested in the subsequent dynamics of the hairpin. Thus, the dynamics of interest is limited to the system

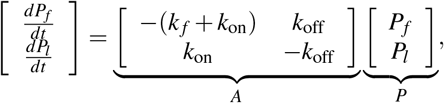

with 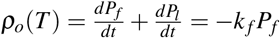

The matrix *A* has two negative eigenvalues *λ*_+−_ that verify :

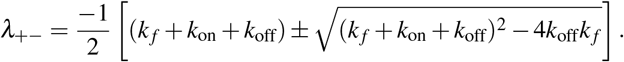

Two corresponding eigenvectors are :

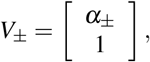

with

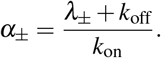

We deduce the two combinations of *P*_*f*_ and *P*_*l*_ whose decay is exponential :

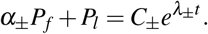

The integration constants *α*_±_ can be deduced from the initial conditions (*P*_*f*_ (0) = 1, *P*_*l*_ (0) = 0), resulting in :

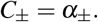

We deduce the expression for *P*_*f*_ (*t*) :

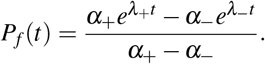

And the expression for the distribution of times spent in the open states :

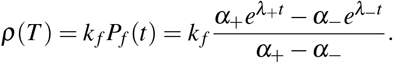

The mean time spent in the open states 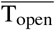 reads :

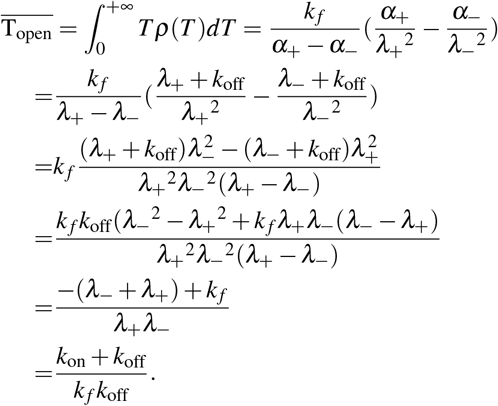

where we used that *λ*_+_ + *λ*_−_ = −(*k*_on_ + *k*_off_) and *λ*_+_*λ*_−_ = *k*_*f*_*k*_off_

*k*_on_ is proportional to the concentration of oligonucleotides : 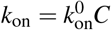. Considering that the energy of hybridization verifies : 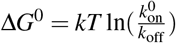, we get:

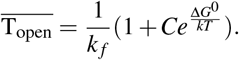

Fitting 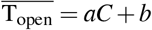 as a linear function of the concentration thus allows us to retrieve 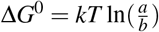

### S.II Resolution in the limiting case: *k*_off_ ≪ *k*_*f*_ and *k*_on_ ≪ *k*_*f*_

This is the case where the kinetics of binding/unbinding is much slower that the kinetics of the fluctuations of the hairpin. This is the case for the binding of the 7nt-oligo and for the case of RecQ binding.

Developing at the order 2 in 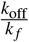 and 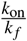, this gives

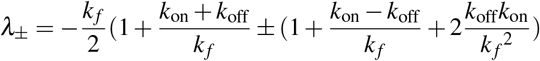

The first order is sufficient to deduce the limit of *λ*_±_. We find :

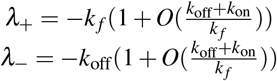

In this limit, the distribution of times is thus a double exponential with very separated times that can be resolved. The long times correspond to 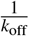 and the short times to 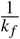.

On the order hand, the proportion of the short times is :

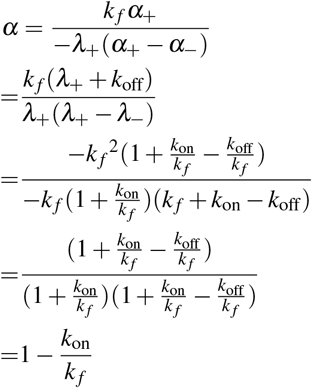

### S.III Resolution in the limiting case: *k*_off_ ≫ *k*_*f*_ and *k*_on_ ≫ *k*_*f*_

This is the case where the kinetics of binding/unbinding is much faster that the kinetics of the fluctuations of the hairpin. This is the case for the binding of short oligonucleotides (5,6 bases).

In this case, the first eigenvalue reads :

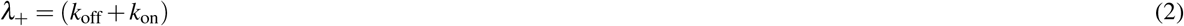

This corresponds to short times that can not be observed.

The second value reads, at the first order in 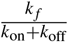 :

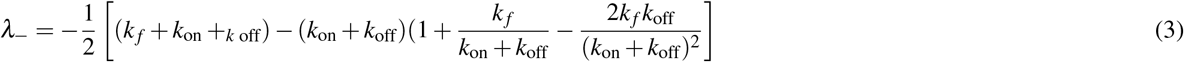

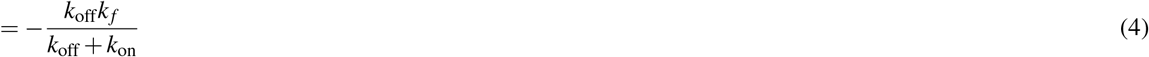

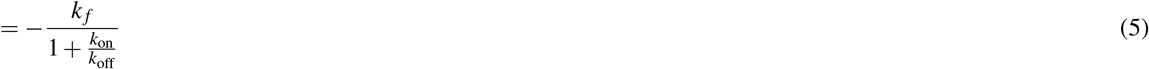

This eigenvalue corresponds to a time that evolves in the same way that the average time 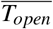, increasing linearly with the concentration. Thus, in this regime, the observable distribution follows a simple exponential distribution that does not allow measuring *k*_on_ and *k*_off_. Only their ratio, and thus Δ*G*, can be measured.

### S.IV DNA sequences

**Table S2.**
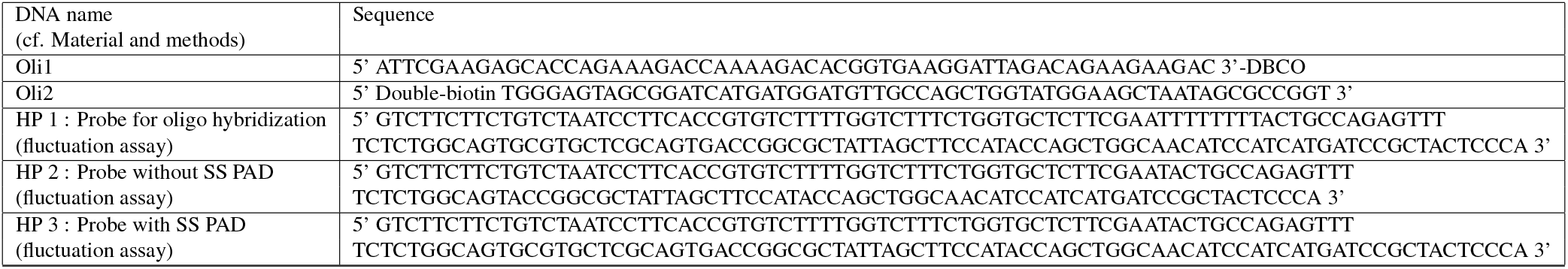
DNA sequences used in this paper. Names are defined in the section Material and Methods.

### S.V Effect of the force on the measure of Δ*G*_binding_

In this section we detail the impact of the force on the estimation of the binding free energy of oligos Δ*G*_binding_. This effect can be calculated if we know the precise force-extension curves of single-stranded DNA *x*_ss_(*F*) and of double-stranded DNA *x*_ds_(*F*). In this case, the variation of free energy reads :

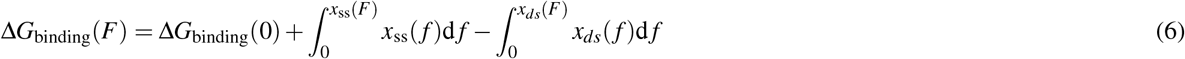

We recall the derivation of this formula for readers that would not be familiar with it in the next section.

The force-extension curve of double-stranded DNA *x*_*ds*_(*f*) is well-established (the worm-like chain model fits it well for forces < 15 pN). The force-extension curve of single-stranded DNA *x*_*ss*_(*f*) is derived from the fit of a long ssDNA by the freely jointed chain model with extension (EFJC, Figure S6). The corresponding estimation of Δ*G*_binding_ as a function of the force is shown on Figure S7. The confidence interval is based on the errors made on the fit of the ssDNA by the EFJC model. The figure shows that the contribution of the force to Δ*G*_binding_ is smaller than 0.04 kcal/mol/nucleotide in the force range of our experiments.

**Figure S6.**
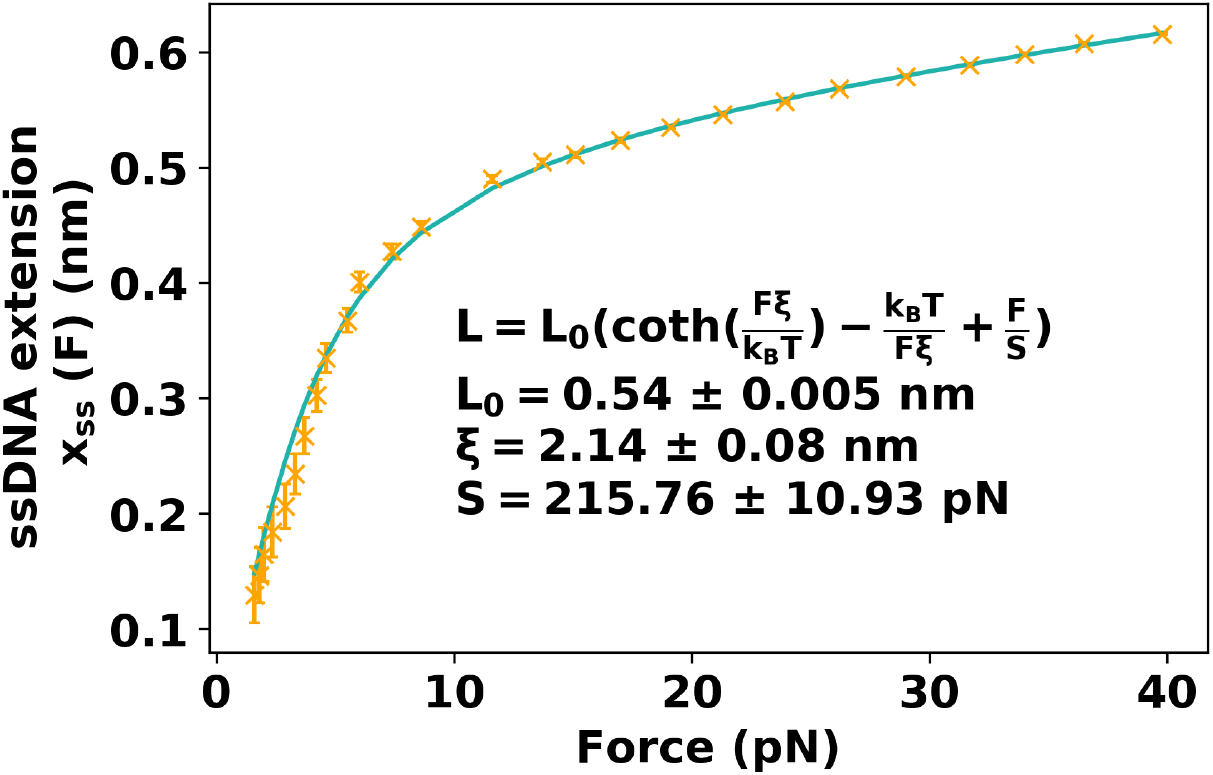
Experimental force-extension curve of a single-stranded DNA molecule and corresponding fit to the freely jointed chain model with extension (EFJC).

**Figure S7.**
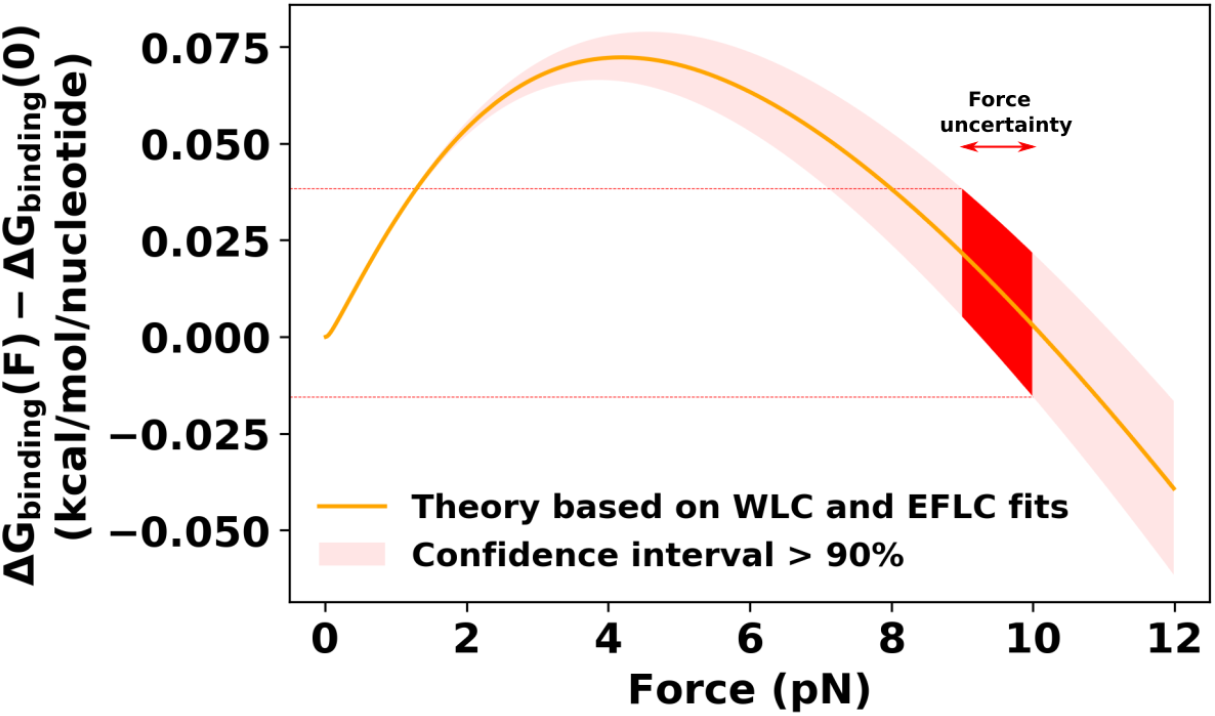
Contribution of the force to Δ*G*_binding_ computed using the worm-like chain model for double-stranded DNA (persistence length of 50 nm and maximum length per nucleotide of 0.34 nm), and the freely jointed chain model with extension for single-stranded DNA.

### S.VI Derivation of the force dependence of Δ*G*_binding_(*F*)

First, we remind how the force-extension curves allow computing the free energy of DNA at various extensions *G*_ss/ds_(*x*_ss/ds_). When pulled at a force F, a molecule adopts an extension that minimizes the free energy of the whole system, consisting of the molecule and the magnetic bead, and that reads :

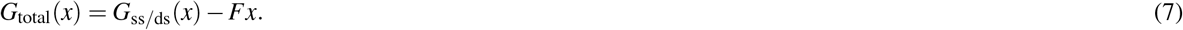

Minimizing *G*_total_ with respect to *x* leads to :

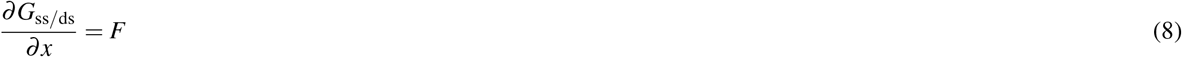

The observed *x*_ss/ds_ is the value of the extension x that minimizes this free energy and is the one measured experimentally. It allows to define a function *x*_ss/ds_(*F*). We can also define the reciprocal function *f*(*x*_ss/ds_) that corresponds to the force F needed to observe the extension *x*_ss/ds_. In particular, it means that we can compute *G*_ss/ds_(*F*) by integrating the observed extension curve with respect to x :

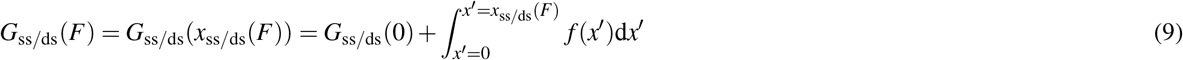

Δ*G*_binding_(*F*) is the difference of free energy between a double-stranded DNA pulled at a force F (final state, hybridized) and two single-stranded DNA, the first being pulled at force F, and the other, free in solution, on which no force is applied :

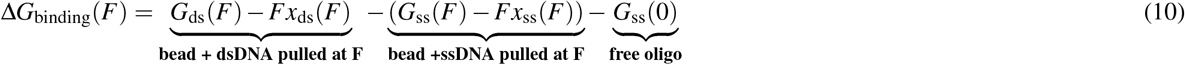

Thus, using the equation just above :

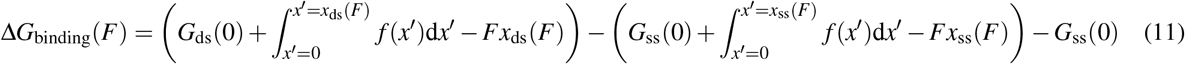

In the two parenthesis, we recognize an integration by part, and thus :

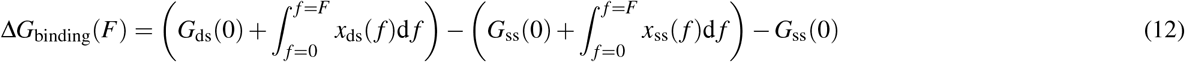

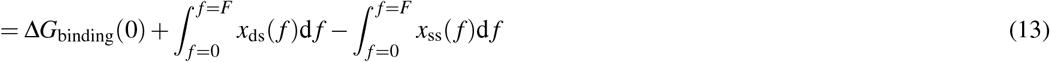

